# Improvement in peripherical visual attentional performance in professional soccer players following a single neurofeedback training

**DOI:** 10.1101/2021.12.13.472447

**Authors:** Sacha Assadourian, Antony Branco Lopes, Arnaud Saj

## Abstract

The effectiveness of EEG-neurofeedback (EEG-NFB) in modulating cognition has been the subject of much research for several years, particularly in relation to attentional functions in healthy subjects and those with attentional deficits. However, its effectiveness on sports performance remains poorly studied and its use is not widely practised among athletes, notably because of its accessibility and questionable effectiveness. The aim of this study is to show that this technology can be accessible, and that Alpha EEG-NFB is immediately effective. Fifteen professional soccer players took part in this study. Using a novel EEG headset that can be installed in less than one minute, and new processing software, the players performed two peripherical attentional tasks before and after, immediately and one month, a single Alpha EEG-NFB training session. The results showed a significant effect on both tasks immediately after EEG-NFB training, with a benefit of more than 30% and this performance continued after one month (20%). This study, the first to use this headset and software, shows that the improvement in sports performance can be related to cognitive performance, especially peripherical visual attentional functions. Furthermore, it demonstrates that the use of the EEG-NFB is accessible and effective for high-level athletes.

## Introduction

There have been many innovations in the field of performance enhancement for athletes, but these have mainly been aimed at improving the physiological, biomechanical, nutritional, and psychological (Mujika et al., 2018) aspects of performance. This physical preparation was then coupled with a mental preparation of the athletes, but sports performance is multifactorial, with notable cognitive aspects (Mujika et al., 2018) but also cerebral. Taking the latter into account is the challenge of the years to come with the development of cerebral preparation of athletes’ performance. The objective of this article is to contribute to the implementation of cerebral preparation by proposing an innovative training procedure dedicated to the increase of the attentional performance of athletes, by tackling the two main challenges: accessibility and efficiency.

The first challenge concerns the accessibility of data processing and the implementation of cerebral preparation through consistent training (Gong et al., 2021). Electroencephalography (EEG) seems to be the most suitable tool, as it is relatively cheap, transportable, and has a very good temporal resolution. Although the EEG is mainly used in clinical medicine, its use has not been widely popularised in sports science. There are many reasons for this: long preparation time before training and duration of training; installing the EEG cap; applying electrode gel; adjusting electrode impedance, etc.

The second challenge is to design effective training protocols for the cognitive abilities targeted by athletes. For football (soccer) practice, the central and peripheral attention in the dynamic visual environment is crucial. Indeed, football players need to be able to mobilise their attention from a spatial and visual point of view. This cognitive ability is possible through the Selective Visuo-Spatial Attention (SVSA). The SVSA can be defined as the ability of committing attention to a position located in our central or peripheral field of view without any overt eye movement (Posner, 1980; Pierce & Saj, 2019). For example, when a player wants to pass the ball to his teammate without telegraphing his intention to the opponents, the player should not look at his teammate and hastily take the decision. Instead, the player must gather information from the environment and stay focused without revealing his intentions to avoid the defensive actions of his opponents. To accomplish this, he must use his SVSA. Thus, a player who has a better SVSA ability, can notice his teammate earlier and make a pass with a higher success rate. For football goalkeepers, high SVSA abilities are essential because they must follow the movements and positions of the players on the field as well as the ball carrier.

In the literature, an increasing number of studies have demonstrated that the EEG alpha activity is closely linked to SVSA, notably by a lateralised modulation of α-power (8–14 Hz) in the parietal (Foxe and Snyder, 2011; Rihs et al., 2009) and occipital (Jeunet et al., 2020; Schmidt et al., 2010) cortices. More recently, EEG Alpha activity (i.e. the SVSA) has been modulated with neurofeedback (NFB) training. Specifically, alpha-synchronisation plays a causal role in the modulation of attention and visual processing (Bagherzadeh et al., 2020). NFB refers to an operant conditioning paradigm in which the individual learns to self-regulate the electrical activity of his/her brain. In EEG-based NFB, EEG is recorded from one or more electrodes placed on the scalp and the relevant components are extracted and fed back using an online feedback loop in the form of audio, visual or combined audio-visual information (Ros et al., 2020). This technique has shown positive effects on the improvement of SVSA and success in sport performance (Gong et al., 2021), and the treatment of many neurological disorders (for example, Saj et al., 2021; Deiber et al., 2021). Taken together, the evidence demonstrates that people indeed obtain benefits through altering the specific EEG activity through NFB.

Therefore, the current study has two aims: the first is to make the use of EEG-NFB in sports practice accessible; the second is to propose an effective protocol using EEG-NFB as a neuromodulatory tool to examine behavioural performance in the central and peripheral SVSA in professional football players using a single session, within-subject design.

## Method

### Participants

Fifteen professional soccer players (15 men; aged 17.6 ± 1.0 year-old) in the Racing Club of Lens (RC Lens®) took part in this one session experiment. The level of performance is divided between U17 and N2: 2 in U17; 5 in U19; 8 in N2. The repartition of field positions was: 5 strikers; 4 midfielders; 3 defenders; 3 goalkeepers. All participants were righthanded. Prior to participating in the study, a consent form was signed and completed. The experimental procedure was led in accordance with the Declaration of Helsinki and was approved by the University of Montreal Ethics Committee (#CEREP-21-037-P).

### Performance Measurement

The set of behavioural measures consists of two attentional tests: an attentional maintenance and distraction resistance task, and a peripheral visual field task. Subjects were seated in a comfortable chair with adjustable height to keep their eyes centred on the screen, located 53 cm away.

#### Attentional maintenance and distraction resistance task

The attentional maintenance and distraction resistance task (Robineau et al., 2014) involves the ability to perceive stimuli simultaneously as quickly as possible (Figure 1A). It is designed to measure visual extinction during simultaneous stimulation at two locations in the visual field. Gabor stimuli will be presented either with an eccentricity of 10° to the left of the fixation cross, 10° to the right, 10° to both sides, or without stimuli. The thresholds range from 1% to 100% contrast (1, 5, 10, 20, 30, 40, 50, 60, 70, 80, 90 and 100%). The participant is asked to indicate whether he or she has seen one of these four possibilities (right, left, both, none) using a keyboard. The Gabor stimuli, presented for a duration of 500 ms, appear on the left, right, both sides of the screen or not in a randomised trial order. Auditory cues (duration: 50 ms; frequency: 1000 Hz) presented simultaneously with the visual stimuli will instruct participants to respond as accurately and as quickly as possible. Participants fixated on a central fixation cross throughout the trials. The number of correct responses is only considered during the 500ms of stimulus presentation. No feedback regarding to the accuracy of the given responses is given to participants.

**Figure 1.**
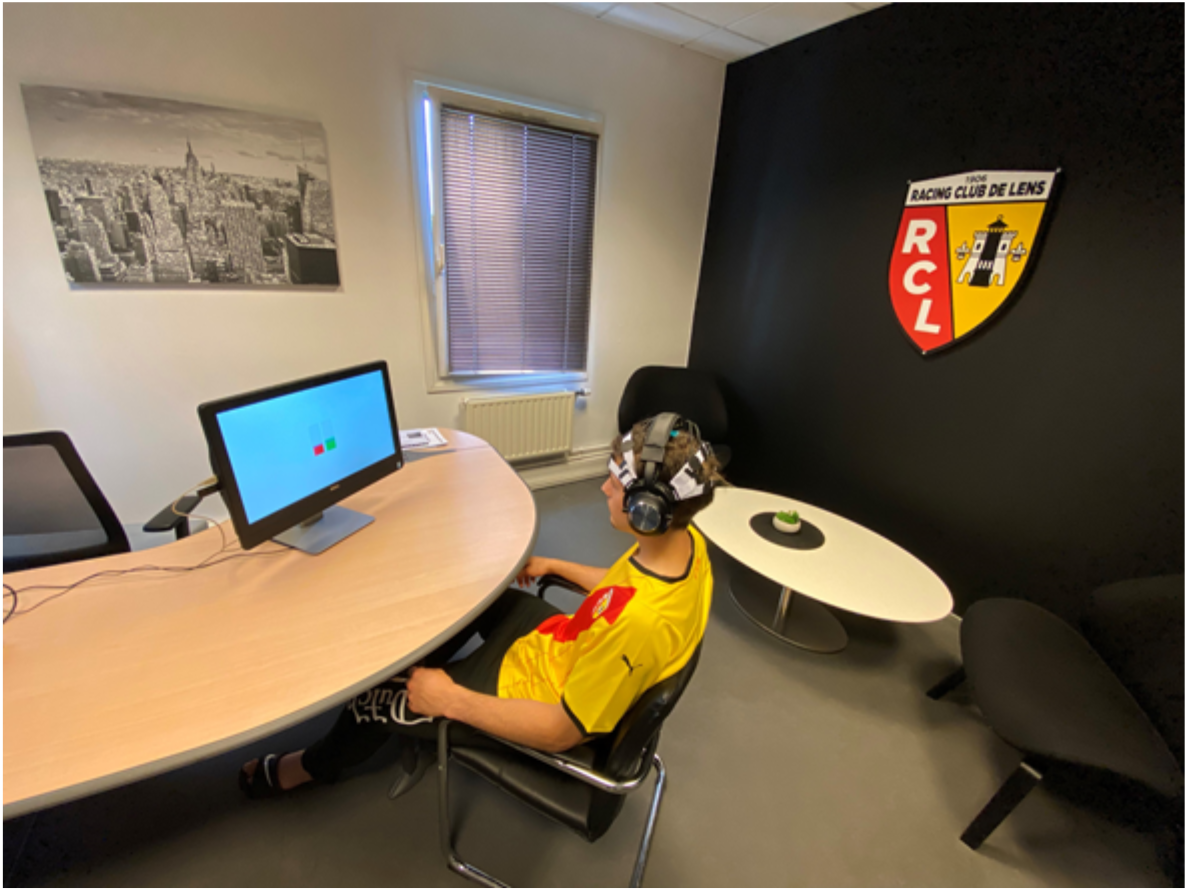
Spectre Biotech headset and software in a football player.

#### Peripheral visual field task

The peripheral visual field task is a modified version of a previously used adaptation (for more details, Nan et al 2014). It consists of reacting as quickly as possible to the appearance of 3 stimuli that are identical either in geometric shape or in colour (Figure 1B). It is designed to measure dynamic peripheral visual capacity. Five objects are presented at the four corners and in the centre of the screen. Each object is positioned in a 6.5 cm diagonal square (placement cue). The objects can take four shapes (circles, horizontal stripes, triangles and vertical stripes) of seven different colours (black, blue, brown, green, red, white and yellow). The test is a target if three of the five objects are of the same colour and shape; otherwise, it is a non-target. Each test session consists of 56 trials, 14 target and 42 non-target trials, with all combinations of different colours and shapes presented. When participants perceive an object combination that forms a target trial, they must press the space bar as quickly as possible. The number of correct responses is only considered during the 500ms of stimulus presentation. No feedback regarding to the accuracy of the given responses is given to participants.

### EEG Neurofeedback Training

The EEG neurofeedback (EEG-NFB) training session consisted of seven blocks of 3 min with 1 min of rest in between each block. During each block, a computer running Spectre-Biotech^®^ software program extracted the signal from each lead and simultaneously calculated the alpha frequency power using a fast Fourier transform algorithm with Hanning windowing function. The signal was 8–13 Hz band-pass filtered using the 6th order Butterworth IIR filter and averaged continuously every 5 ms. The resulting values were then displayed to participants on-screen via bar charts displaying alpha power at the bilateral parietal cortex, corresponding to the P3 and P4 electrode in the 10–20 international EEG positioning system, and an auditory tone if the threshold was exceeded.

### Procedure

An EEG headset developed by Conscious Labs^®^ in partnership with Spectre Biotech® was placed on the participants’ heads (Figure 2). The EEG Headset is a 16-Channel EEG SUPRA Headphones wireless and is implemented in a few minutes only. The 16-channel are dry and flexible electrodes (dry active ThinkPulseTM polymer electrodes: replaceable, flexible, biocompatible & multiple shapes). The connectivity is done by BLE wireless transmission (Bluetooth Low Energy 4.2) via USB dongle. The participants were seated in a quiet room in front of a screen. Each participant performed the two behavioural tasks before (t0) and immediately (t1) and one month (t2) after the EEG-NFB training. No adverse effects or unusual symptoms were reported by any participant either before, during, or after the EEG-NFB training sessions.

**Figure 2.**
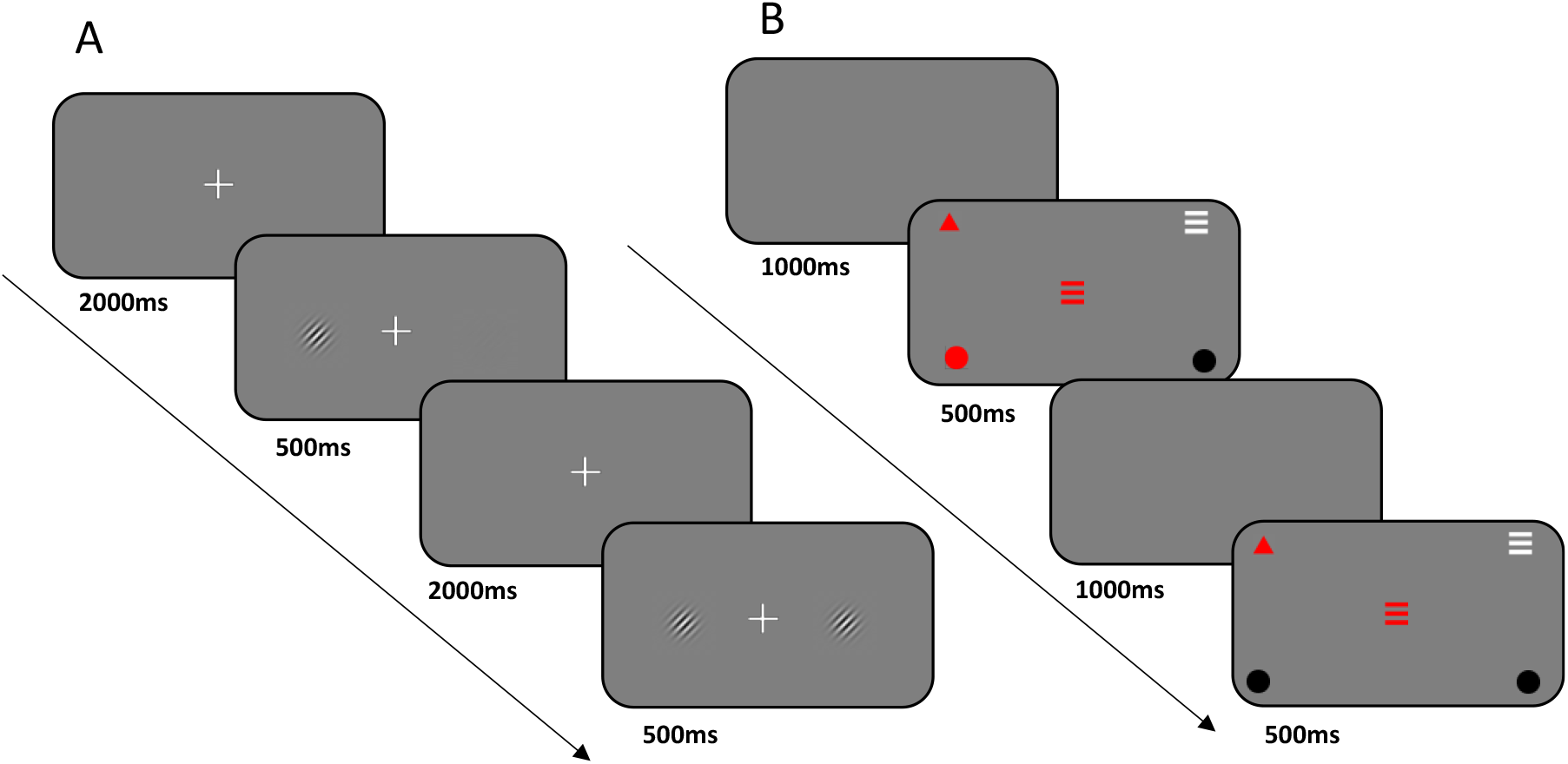
Example of trail in the peripherical visual attentional maintenance and distraction resistance task (panel A) and in the Peripheral visual field task (panel B).

## Data analysis

Data analysis consisted of comparing subjects’ performances (correct responses) on the two tasks before (t0) and after (t1 and t2) a single session of EEG-NFB. First, a Shapiro-Wilk normality test was performed to show whether the data were normally distributed. Secondly, a Student’s t-test on the before-and-after variables as a function of the task was performed, as well as measuring the effect size with Cohen’s d. The significance level was set at p<0.05.

## Results

The single EEG-NFB session has a significant effect on both behavioural tasks (Figure 3). Normality test (Shapiro-Wilk) showed that the data were normally distributed, for the attentional task (p=0.240) and visual field (p=0.573). Participants significantly improved their performance immediately after the EEG-NFB session, going from 18.5%+9.9 correct responses to over 30.3%+12.9 (t(14) = 3.69, p < 0.01; Cohen’s d =0.95) in the Attentional maintenance and distraction resistance task and from 7.5%+4.3 to over 34.5%+14.5 (t(14) = 7.85, p < 0.01; Cohen’s d (2.03) in the Peripheral visual field task. After one month, performance (t2=24.7%+9.9) was still significantly better than at t0 (t(14) = 2.29, p < 0.01; Cohen’s d =1.35), but it also lost power compared to t1 (t(14) = 4.91, p < 0.01; Cohen’s d =0.93).

**Figure 3.**
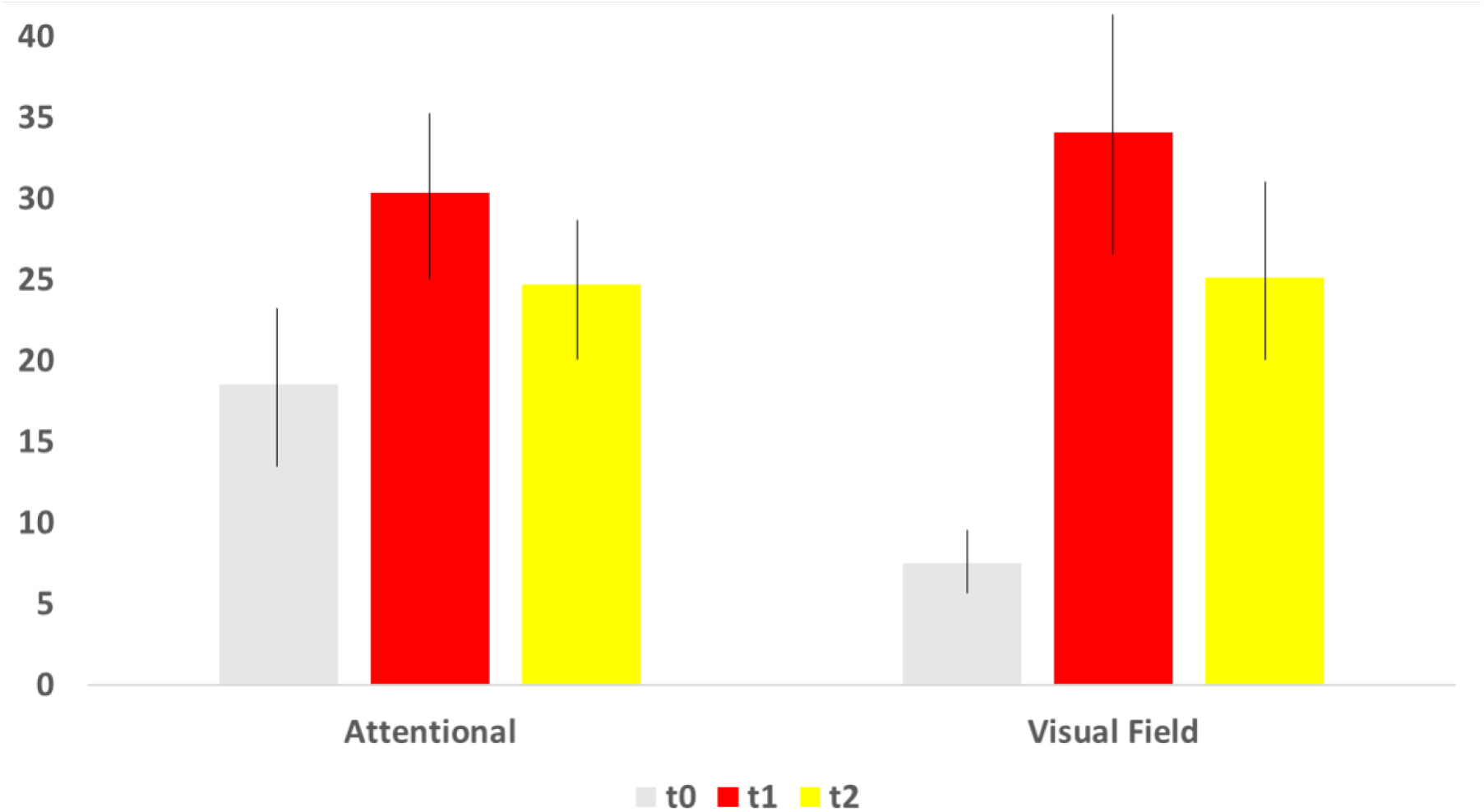
Correct responses t0, t1 and t2 EEG-NFB training for the peripherical attention and visual field task.

## Discussion

The objectives of this study were to investigate accessibility and efficiency of the effect of Alpha neurofeedback training on football players’ performance. Concerning the accessibility, the set-up time for the headset and training took no more than 5 minutes. The use of dry electrodes has proven to be close to the effectiveness of traditional electrodes (Collado-Mateo et al., 2015). The second aim was the efficiency of EEG-NFB training. Our results showed that football players receiving Alpha neurofeedback training demonstrated enhanced performance in two attentional tasks just after one session, compared with the initial performance. This performance remains a month training. This finding lends preliminary support to the hypothesis that Alpha EEG-NFB is effective for reducing Alpha power and leads to superior football (soccer) performance.

This result confirms the results of previous studies on the improvement of performance in football players (Wenya et al., 2013; Wenya et al., 2014) or sportsmen in general (see Gong et al., 2021). The major element in this study is that performance is increased with only one session of Alpha EEG-NFB. Indeed, most studies suggest 2 to 5 weeks of daily training to see effects on cognition, mental health, and performance (Nan et al., 2013; Nan et al., 2014; Noortje et al., 2016). The only studies with a single session are studies in patients with attentional disorders (Saj et al., 2018; Deiber et al., 2020). The first study (Saj et al., 2018) involved brain damaged patients with an attentional disorder: spatial neglect (see Vuilleumier et Saj, 2013). This disorder is characterised by an over-focused attention with a visual-spatial bias. In other words, patients appear to be attracted to a target, like a magnet is to steel. In their study, the authors were able to correct this bias in a single session, allowing the patients to have a more diffuse attention. The second (Deiber et al., 2020) concerns patients with ADHD. This disorder is characterised by an overly diffuse attention, with a difficulty to focus on a single target for a long period of time. Here too, a single session was used to correct this bias to make it less diffuse. These two studies are seminal works for understanding what we have been able to highlight in our study. Indeed, our Alpha EEG-NFB session allowed subjects to better focus their attention on the targets but also to make it more diffuse by fetching all the available information. These two types of phenomena are essential to the good reading of the game during a football match.

Our study has two limitations but also two perspectives. The first limitation is that this increase in attentional performance should be observed in a more ecological environment, i.e. on the football field. The second limitation concerns the maintenance of this increase in performance over time. How long does this increase last? When should another session be offered?

The first perspective is that a single session allows players to improve their performance but also to return to their initial levels in case of a drop in performance, for example at midseason or when external factors reducing performance occur such as fatigue, stress, poor pressure management or downtime after injury, etc. The second perspective is that our technology will provide a solution to the perceived difficulty in using EEG-NFB headset in sport (Gong et al., 2021). Indeed, the spectre biotech® equipment used is quick to install and use, with the same performance as traditional headsets but without the constraints. The third perspective is more clinical. Indeed, the accessibility of the tool opens new possibilities in diagnostic assistance (Newson and Thiagarajan, 2019) and therapeutic management (Saj et al., 2021).

***In conclusion***, a single Alpha EEG-NFB training can increase the peripherical visual attentional performance of professional football players using a simple and fast tool. The headset is placed on the athlete’s head in less than a minute. He/she only has to connect and follow the instructions of the software. The program has been developed to outsource all EEG processing. There is no need to be trained, making this tool accessible to the club’s medical team and easily used by a health professional, with an easy and accessible interface. Athletes can therefore set up their own training programs and follow them anywhere and at any time during the season.

## Acknowledgements

The authors would like to thank the Racing Club de Lens, in particular the head of the clubs academy, Mr Eric Assadourian, for his welcoming and the space, as well as all the soccer players who took part in this study.

